# A minimal model of protein allocation during phototrophic growth

**DOI:** 10.1101/183236

**Authors:** Marjan Faizi, Tomáš Zavřel, Cristina Loureiro, Jan Cerveny, Ralf Steuer

## Abstract

Photoautotrophic growth depends upon an optimal allocation of finite cellular resources to diverse intracellular processes. Commitment of a certain mass fraction of the proteome to a specific cellular function, typically reduces the proteome available for other cellular functions. Here, we develop a minimal semi-quantitative kinetic model of cyanobacterial phototrophic growth to describe such trade-offs of cellular protein allocation. The model is based on coarse-grained descriptions of key cellular processes, in particular carbon uptake, metabolism, photosynthesis, and protein translation. The model is parametrized using literature data and experimentally obtained growth curves. Of particular interest are the resulting cyanobacterial growth laws as fundamental characteristics of cellular growth. We show that the model gives rise to similar growth laws as observed for heterotrophic organisms, with several important differences due to the distinction between light energy and carbon uptake. We discuss recent experimental data supporting the model results and show that minimal growth models have implications for our understanding of the limits of phototrophic growth and bridge a gap between molecular physiology and ecology.

## 1 Introduction

Phototrophic microorganisms, including cyanobacteria, have recently gained significant interest as potential host organisms for the renewable synthesis of pharmaceuticals, food supplements, and chemical bulk products, including biofuels [1, 2]. As the only known oxygen-evolving photoautotrophic prokaryotes, cyanobacteria posses several advantageous properties that make them promising candidates for biotechnological applications: cyanobacteria use sunlight, water, and atmospheric CO_2_ as their primary source of energy, reducing power, and carbon, respectively. They thrive in almost all environments, from Antarctica to hot springs and deserts, as well as from fresh water to high saline ecosystems [3]. Cultivation is possible in salt and brackish water, thereby avoiding direct competition with conventional agriculture. Cyanobacteria are genetically tractable and exhibit high photoautotrophic growth rates.

The latter, the capability to grow with high rates in diverse environments, is frequently emphasized as a key requirement for applications in green biotechnology [4, 5, 6]. Many cyanobacteria, including well-characterized model strains, have typical doubling times of 7-12 hours (h) or more [6, 4]. The fastest reported doubling time of a cyanobacterium to date is 1.9h [5]. Given the importance of fast growth as an indicator of overall culture productivity, however, the (molecular) limits of phototrophic growth are still insufficiently understood. Only recently, a number of computational [4, 7, 8, 9], as well as experimental [6, 5] studies have begun to investigate the molecular limits of phototrophic growth. In this work, we develop a minimal kinetic model of cyanobacterial phototrophic growth based on a coarse-grained description of relevant intracellular processes. Using the computational model, we seek to understand the organization of phototrophic growth in terms of the cellular ‘protein economy’ [10, 4, 7, 9] of growth. That is, we seek to understand how a growing cell may optimally allocate its limited energy and protein resources to different intracellular processes relevant for phototrophic growth, including protein translation, photosynthetic electron transport, carbon uptake and metabolism.

Our approach is based on similar models already available for heterotrophic growth. Since the seminal studies of Jacques Monod describing the growth of a bacterial culture [11], a wealth of information has been acquired with respect to the physiology of growing bacterial cells [12, 13]. Of particular interest are fundamental characteristics of growth that are independent of the chemical nature of the medium, such as the gross chemical composition in terms of protein, RNA, DNA, carbohydrates and lipids [13]. An early key observation was that several of these characteristics are simple monotonic functions of growth rate [14, 13] - in particular the concentration of ribosomes was found to be a linear function of growth rate [15].

Since then, a number of experimental and theoretical studies have addressed the covariation between the cellular composition of macromolecules and the growth rate for heterotrophic microorganisms [14, 10, 16, 17, 18, 19, 20]. These models are minimal whole-cell models that describe fundamental processes of cellular growth by coarse-graining the proteome into few essential classes. Properties of the models are then typically evaluated in the context of evolutionary *optimality*: the allocation of proteins to the respective processes is assumed to be optimal in the sense that the growth rate in a given environment is maximal, and that synthesizing more protein within one class, at the expense of other classes, would lower the overall growth rate [14]. While there are certainly caveats concerning the assumption of optimality, this assumption has proven to be a useful starting point to investigate and benchmark protein allocation in growing cells.

With the exception of the recent study of Burnap [4], however, similar models have not yet been developed and analyzed for phototrophic growth. We therefore propose a minimal kinetic model that allows us to describe the optimality of the macromolecule composition for cyanobacterial growth under different environmental conditions. The model is based on available resource allocation models for heterotrophic growth, in particular the models of Molenaar et al. [10], Maitra and Dill [19] and Weisse et al. [20], but accounts for the specific properties of photoautotrophic growth. Going beyond the model of Burnap [4], we consider a primitive CO_2_-concentrating mechanism and the consequences of photodamage on protein allocation. The model is parametrized using experimentally determined growth curves for the cyanobacterial strain *Synechocystis* sp. PCC 6803. Our key questions are: (i) does a minimal model of cyanobacterial growth allow us to reproduce experimentally observed growth curves? (ii) does a model of phototrophic growth give rise to similar growth laws as observed for heterotrophic growth? (iii) how do potential photodamage and carbon cycling impact observed growth laws, and (iv) what are the implications of minimal growth models for biotechnology, ecology and our understanding of the limits of phototrophic growth?

## 2 A minimal model of phototrophic growth

To develop a minimal model of phototrophic growth, the relevant cellular processes are coarse-grained into three cellular functions: (i) a minimal carbon metabolism consisting of carbon uptake and anabolic reactions, (ii) photosynthesis that provides cellular energy (and reducing power), and (iii) protein translation. The proteome is represented by four different protein classes: transporters (*T*) that facilitate uptake of inorganic carbon, metabolic enzymes (*M*) that catalyze carbon assimilation and anabolic reactions, ribosomes (*R*) that facilitate protein translation, and photosynthetic proteins (*P*) that produce cellular energy. The model structure is shown in figure 1.

**Figure 1:**
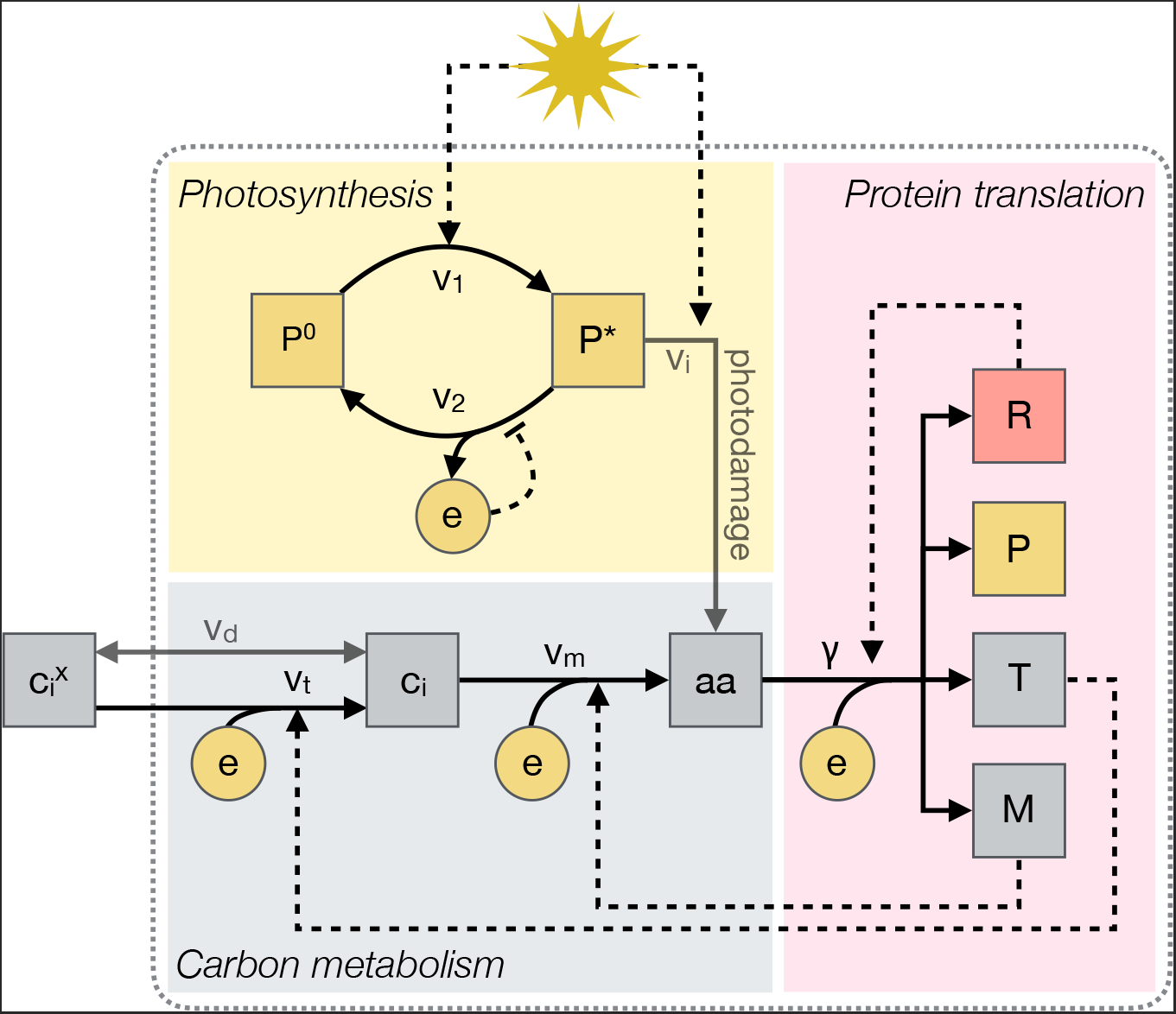
A minimal model of phototrophic growth. External inorganic carbon 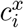 is actively imported into the cell via a transporter *T (v_t_)*. The metabolic enzyme *M* converts intracellular inorganic carbon molecules *c_i_* into amino acids *aa (v_m_)*, the precursors for growth. Protein synthesis (*γ*) is facilitated by ribosomes *R*. All reactions depend on a cellular energy source, denoted as e. Cellular energy is obtained by the photosynthetic light reactions. The photosynthetic unit *P* exists in two states: active (*P*\*) and inactive (*P*^0^). *P*^0^ is activated by light absorption (*v*_1_), and energy is released during the transition between the activated to the inactivated state (*v*_2_). The latter reaction is subject to product inhibition. The model is later extended to account for uptake (and loss) of inorganic carbon by passive diffusion (*v_d_*) and for photodamage (*v_i_*).

The model consists of 7 ordinary differential equations (ODEs) describing the dynamics of all internal compounds in units of numbers of molecules per cell. For simplicity, and following Weisse et al. [20], we assume that the average cell volume is constant. In the initial model, we do not consider passive uptake or loss of inorganic carbon (carbon cycling) and damage induced by excessive light (photoinhibition). In the following, we briefly outline core components of the model, the full system of ODEs is provided in the supplementary text (sections 1–2).

### 2.1 Carbon assimilation and metabolism

Uptake of inorganic carbon and its assembly into a (generic) amino acid *aa* is described in two steps: external inorganic carbon 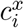 is irreversibly imported (reaction *v_t_*), facilitated by the transporter protein *T*, subsequently, intracellular inorganic carbon *c_i_* is assimilated into organic carbon and converted into the amino acid *aa* (reaction *v_m_*). The reaction *v_m_* is catalyzed by a metabolic protein *M*, which represents all enzymes required to catalyze the conversion from *c_i_* to *aa*. For simplicity, we do not distinguish between CO_2_ and bicarbonate 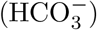. The initial model does not include loss of intracellular inorganic carbon by passive diffusion (carbon cycling). The dynamics of the abundance of intracellular inorganic carbon *c_i_* are governed by the differential equation

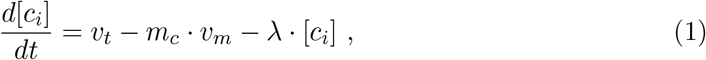

where *m_c_* denotes the number of inorganic carbon molecules *c_i_* required to produce one amino acid. The last term describes dilution by cellular growth, with *λ* denoting the growth rate. Both reaction rates are assumed to follow irreversible Michaelis-Menten kinetics and depend on the abundance of the energy unit *e*,

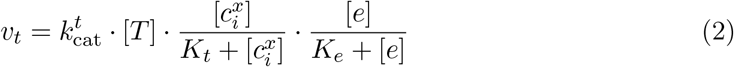

and

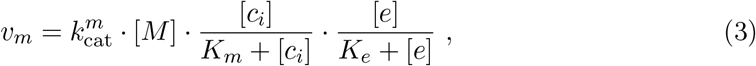

where 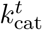 and 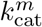 denote the maximal turnover rates of the transporter and metabolic proteins, and *K_t_* and *K_m_* denote the corresponding half-saturation constants. The half-saturation constant *K_m_* of the metabolic protein is set to mimic the low (relative) affinity of the enzyme ribulose-1,5-bisphosphate-carboxylase/-oxygenase (RuBisCO) for its substrate CO_2_. For simplicity, the half-saturation constant *K_e_* is assumed to be identical for all energy-dependent reactions. See section 2.5 for model parametrization.

### 2.2 Ribosomes and protein translation

Protein translation is described analogous to earlier models for heterotrophic growth [10, 19, 20]. Different from Weisse et al. [20], we do not explicitly represent transcription and synthesis of mRNA. Protein complexes are translated by ribosomes *R*, using the precursor *aa* and energy. The translation rate *γ_j_* of a protein complex *j* with a length of *n_j_* (in units of amino acids) is

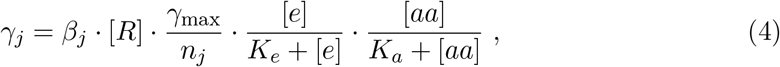

where γ_max_ denotes the maximal elongation rate that is divided by the size of the protein complex *n_j_* to account for the fact that larger complexes take longer to translate [20]. The parameter *K_a_* denotes the half-saturation constant of ribosomes for amino acids and the factor *β_j_* denotes the fraction of total ribosomes allocated to the translation of protein complex *j*. The factors *β_j_* therefore determine the allocation of resources to cellular functions and fulfill the constraint

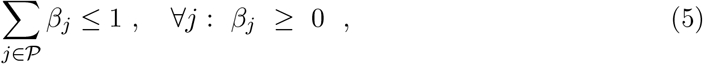

with 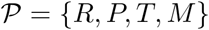. The dynamics of the abundance of each protein complex is given by the difference between translation rate and dilution by cell growth. For example, the amount of ribosomes is governed by the following differential equation (and analogously for all other protein complexes)

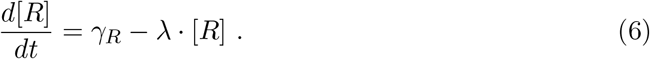

We assume that for fast growing cells, protein degradation is negligible compared to the dilution term describing cell growth (with the exception of photodamage discussed below).

### 2.3 Photosynthesis and the electron transport chain

The conversion of light into cellular energy is described analogously to existing three-state models of phototrophic growth [21, 22, 23]. Specifically, we follow the model of Han [24] and describe light harvesting and the electron transport chain as a single process that is facilitated by a photosynthetic unit *P*. The photosynthetic unit is defined as the assembly of light-harvesting complexes, photosystems II and I, and the photosynthetic electron transport chain [24]. The photosynthetic unit *P* exists in two states: inactivated *P*^0^ and activated *P*\*. Absorption of light facilitates the transition to the activated state *P*\* (reaction *v*_1_ in figure 1). The transition from the activated state *P*\* to the inactivated state *P*^0^ then results in the production of a cellular energy unit e (reaction *v*_2_ in figure 1). For simplicity, we do not distinguish between reducing power and chemical energy (ATP): the energy unit *e* is understood as an abstract entity that combines contributions from ATP, GTP and NADPH.

The photosynthetic cycle is assumed to be fast, compared to the timescales of translation. We obtain an expression that describes the reaction rate *v*_2_ in terms of total photosynthetic unit [*P*] = [*P*^0^] + [*P*\*],

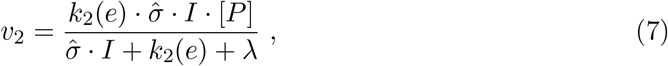

where *k*_2_(*e*) denotes the effective maximal turnover rate of the photosynthetic unit, subject to product inhibition exerted by the energy unit *e*. Light absorption is given by the product of the light intensity *I* and the effective absorption cross-section 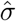 per photosynthetic unit. We note the two limiting regimes for the photosynthetic light reactions are: for high light (lim *I* → ∞), the rate approaches *v*_2_ = *k*_2_(*e*) · [*P*], thus the rate is only dependent on the amount of *P* per cell; for low light 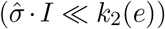, and hence slow growth (λ ≪ 1), the rate approaches 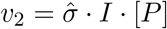 and depends linearly on the absorbed light energy. We assume that per cycle *m_ϕ_* units of *e* are released, hence the total synthesis rate of *e* is *m_ϕ_ · v*_2_. Likewise, we assume that the release of molecular oxygen *O*_2_ is proportional to *v*_2_.

### 2.4 Describing cellular growth

The aim of our study is to investigate cellular growth as a function of protein allocation expressed by the factors *β_j_*. To this end, we require an expression of the growth rate λ as a function of kinetic parameters. Following Weisse et al. [20], we consider the (average) cellular density 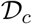 (or mass per volume) to be constant under different growth conditions and for different rates - a fact that is supported by experimental observations [25]. The cellular density 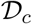 is proportional to the weighted sum of the abundances of cellular components (the mass of components is measured in units of *aa*)

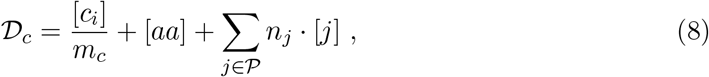

with 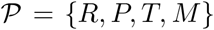. Applying the steady-state assumption on the expression for 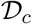 (see also supplementary text, section 2.4), we obtain an expression for the growth rate λ,

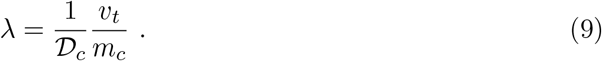

In the following, equation (9) serves as the objective function for the optimization problem. We seek to maximize *λ* as a function of protein allocation, as determined by the factors *β_j_* (the allocation of translational capacity), subject to the constraint specified in equation (5).

### 2.5 Parameterizing the model

We aim for a semi-quantitative model. That is, all relevant parameters should be within reasonable ranges and have a sound justification based on the biochemical literature. Model results, however, are understood as approximations, suitable to investigate general properties of phototrophic growth. Kinetic parameters were sourced from the literature and are summarized in table 1.

**Table 1:**
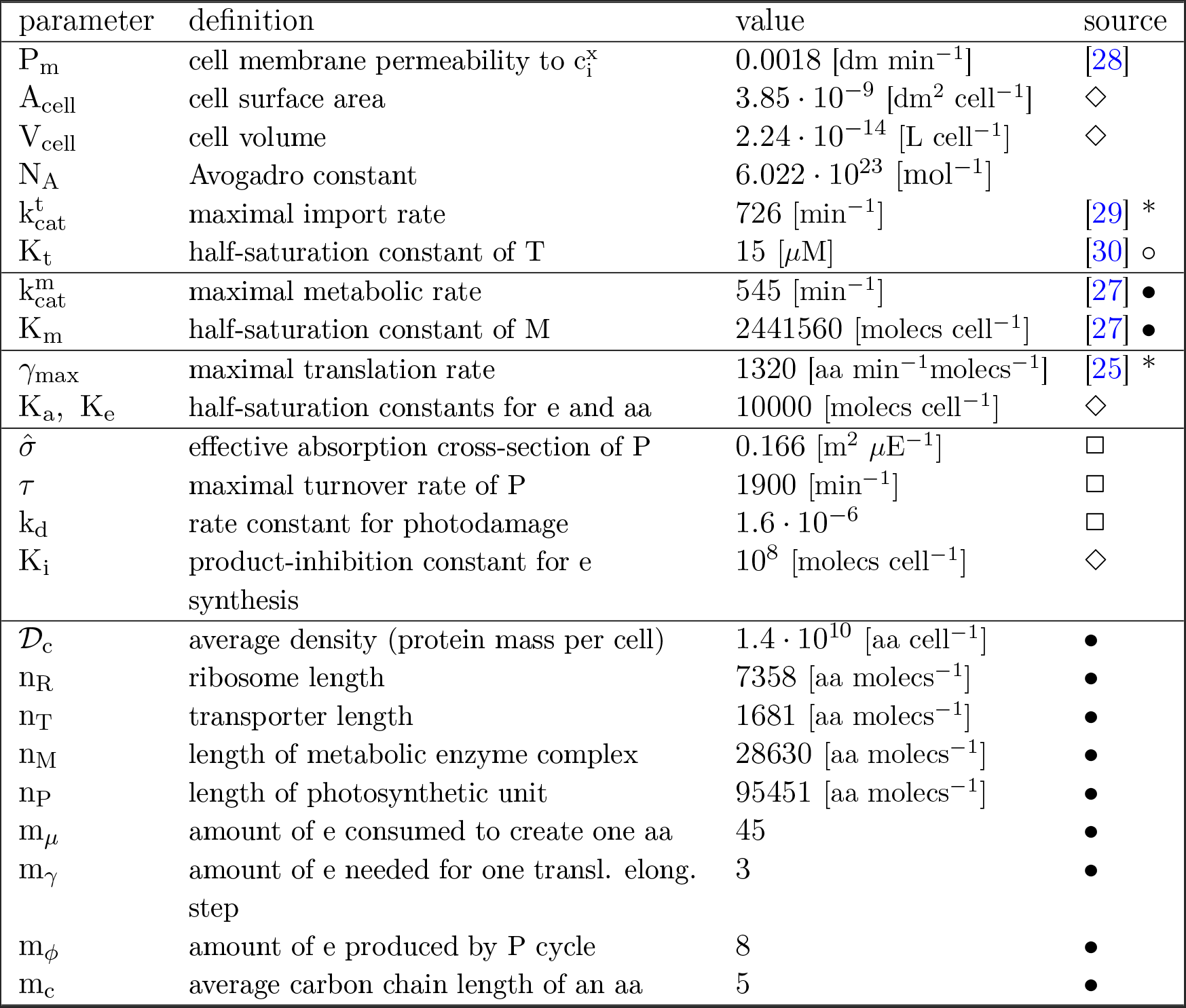
Parameters of the model. Parameters mainly relate to the cyanobacterial species *Syuechocystis* sp. PCC 6803^•^ (and *Syuechococcus* sp. strain PCC 7942°). If no data were available in the literature, parameters are adopted from *E.coli*\*. The remaining parameters are estimated here^⋄^ or fitted^□^ (see supplementary text, section 3.3). Amino acids are abbreviated as *aa* and molecules as *molecs* for the units.

Protein lengths (in units of *aa)* for the macromolecules *R, P*, and *T* were derived using the known molecular composition of macromolecules involved in the respective processes. The respective tables are provided in the supplementary text, tables T2-T9. For the metabolic protein *M*, participating proteins were derived from the metabolic reconstruction of Knoop et al. [26]. Stoichiometric coefficients were approximated as follows: translation is assumed to require 3 energy units (one ATP and two GTP) per amino acid, each photosynthetic cycle results in 8 energy units (joint contributions from ATP and NADPH), and the amount of energy units required to synthesize one generic amino acid *aa* was approximated using the reconstruction of Knoop et al. [26]. The average cell density 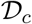 (protein mass per cell in units of *aa)* is calculated by assuming that the mass fraction of proteins considered in the model is ≈ 25% of the cell dry weight (we assume that the total proteome is about 50% of cell dry weight). We do not consider constituent proteins and other cellular components explicitely. Cell surface and volume (*A_ce11_* and *V_cell_*) are calculated by assuming a spherical shape with an average cell radius of 1.75 *μm* (for derivation of the cell radius see supplementary text, section 3.3).

We consider the enzyme ribulose-1,5-bisphosphate-carboxylase/-oxygenase (RuBisCO) as the rate-limiting step in metabolism. Its affinity *K_m_* = 181 *μ*M [27] is converted into molecules per cell using the conversion factor 1 *μ*M = 10^-6^ · *N_A_ · V_cell_* [molecs cell^-1^]. Remaining intracellular half-saturation constants (*K_a_, K_e_*) are set to low arbitrary values. The synthesis of the energy unit *e* is subject to product inhibition using an (arbitrarily set) inhibition constant *K_i_* to prevent unreasonable accumulation of *e*.

Remaining unknown parameters are the effective absorption cross-section 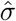 of P, the photodamage rate *k_d_*, as well as the maximal turnover rate of the photosynthetic unit τ, given by

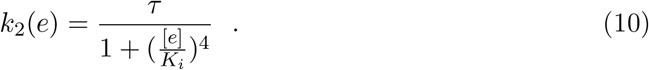

The parameters 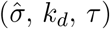 are estimated using experimentally obtained growth curves for *Synechocystis* sp. PCC 6803 under CO_2_-saturating conditions (see section 4.3). The fit was obtained using a reference concentration of external inorganic carbon of 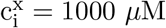. For the initial model the fitted parameters were used, with *k_d_* and *P_m_* equal to zero (neglecting diffusion and photodamage). Details about the estimated and fitted parameters are provided in the supplementary text (section 3).

## 3 The protein economy of phototrophic growth

We seek to evaluate the model with respect to bacterial growth laws. To this end, we vary the proteome allocation, as determined by the factors *β_j_*, to maximize the growth rate λ, specified in equation (9). All computations were carried out in *Python* 2.7 using the package *scipy.optimize*. Parameters are as specified in table 1, except otherwise noted.

### 3.1 Growth kinetics of the minimal model

We solve the optimization problem as a function of the concentration of external inorganic carbon 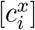 and light intensity *I*. The resulting growth curves, shown in figure 2, are consistent with the Monod equation. Similar results have been obtained for heterotrophic bacteria [10, 20]. The Monod curves allow to estimate the (model-based) maximal growth rate λ^*max*^ under saturating concentrations of external inorganic carbon and light. The resulting value λ^*max*^ = 0.149 h^-1^ corresponds to a doubling time of approximately 4.65h, well within a reasonable range for the growth rate of *Synechocystis* sp. PCC 6803, and slightly slower than the fastest reporting doubling times for cyanobacteria [5, 6].

**Figure 2:**
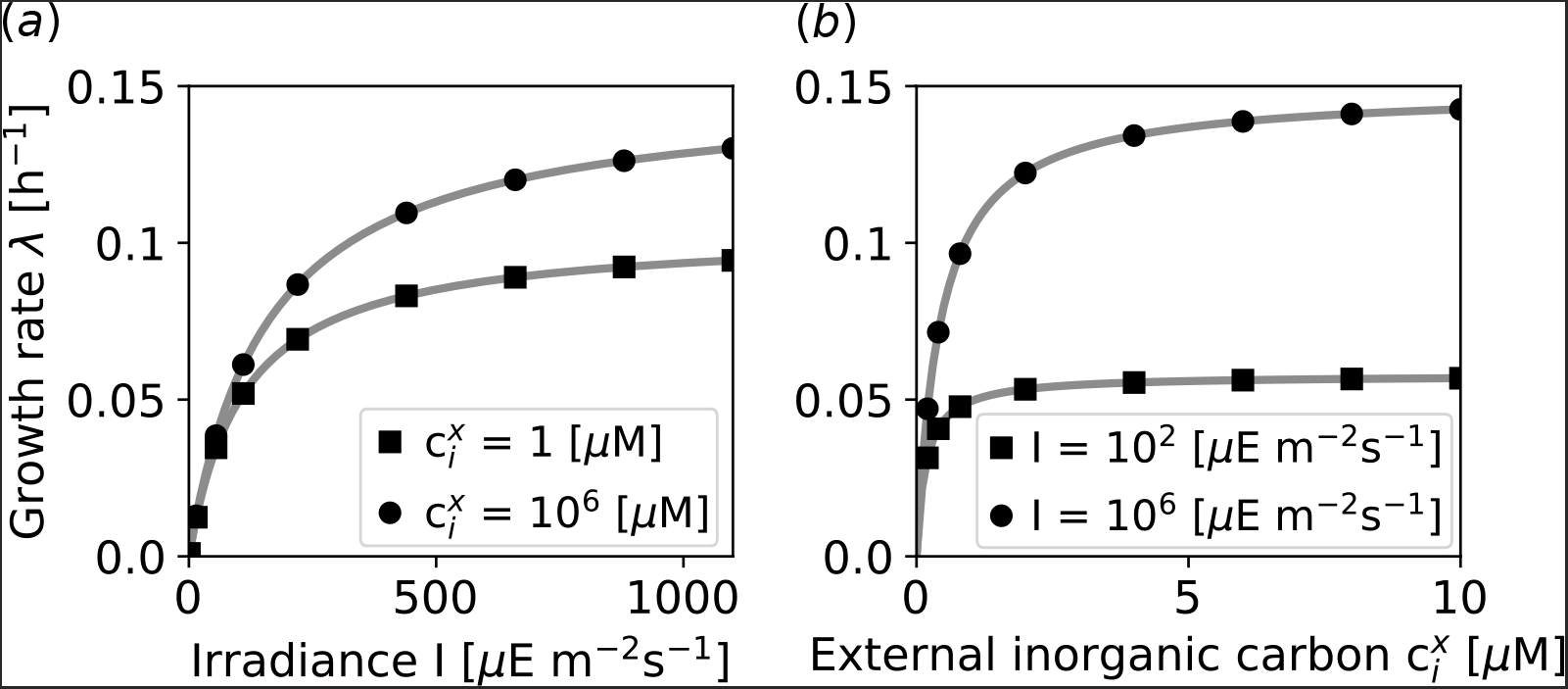
The minimal model reproduces Monod-like growth curves. Maximal growth rates (black circles and squares) are determined by optimizing protein allocation for different values of light intensity *I* and external inorganic carbon 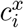. Solid gray lines indicate the Monod equation (11) that was fitted to the model-derived growth rates.

Considering the co-limitation by external inorganic carbon 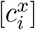 and light availability *I* [31], the growth curve is not consistent with a multiplicative dual-substrate Monod equation. Neither is the growth curve consistent with Liebig’s law of the minimum where only one nutrient is the limiting nutrient. However, a Lineweaver-Burk plot (see supplementary text, figure S3) of growth rate versus light intensity for different inorganic carbon concentrations shows parallel lines, which is indicative of uncompetitive inhibition. Therefore, we conjecture that the growth law is consistent with a rate equation in which absence of a nutrient corresponds to (uncompetitive) inhibition,

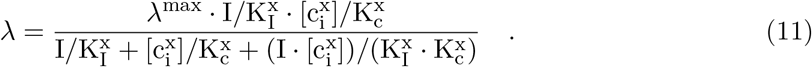

Using equation (11), we obtain the effective half-saturation constants for the external nutrients, 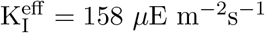 and 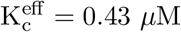, respectively. The half-saturation constant for external inorganic carbon is significantly below the corresponding half-saturation constants for the transporter and metabolic reaction (K_t_ = 15 *μ*M and K_m_ = 181 *μ*M, see section 2.5). The decrease of the effective half-saturation constant for external inorganic carbon indicates that the irreversible transport implements a rudimentary carbon concentrating mechanism (CCM). We observe that for the optimized solution, the intracellular concentration is at all times sufficient to saturate the metabolic enzyme *M*. Likewise, the intracellular concentration of the energy unit e is at all times sufficient to saturate all energy consuming reactions. Saturation of reactions with their respective substrates ensures that intracellular reactions, including protein translation, operate close to their maximal capacity.

### 3.2 Protein allocation and phototrophic growth laws

As the next step, we study protein allocation, optimized for maximal growth rate, as a function of environmental conditions. Figure 3 shows the model-derived optimal protein allocation for phototrophic growth. Using light intensity *I* and external inorganic carbon 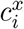 as parameters, the factors *β_j_* were optimized to give rise to a maximal growth rate λ. Corresponding to results obtained for earlier models of heterotrophic growth, the ribosomal mass fraction is a linear function of the growth rate, irrespective of whether *I* and 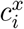 is changed. Indeed, the linear dependence is a direct consequence of the model definitions. Re-arranging equation (6) and assuming that translation is saturated with respect to its substrates *aa* and *e*, we obtain

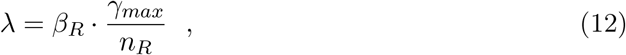

whereby *β_R_* corresponds to the amount of ribosomes allocated to their own synthesis.

**Figure 3:**
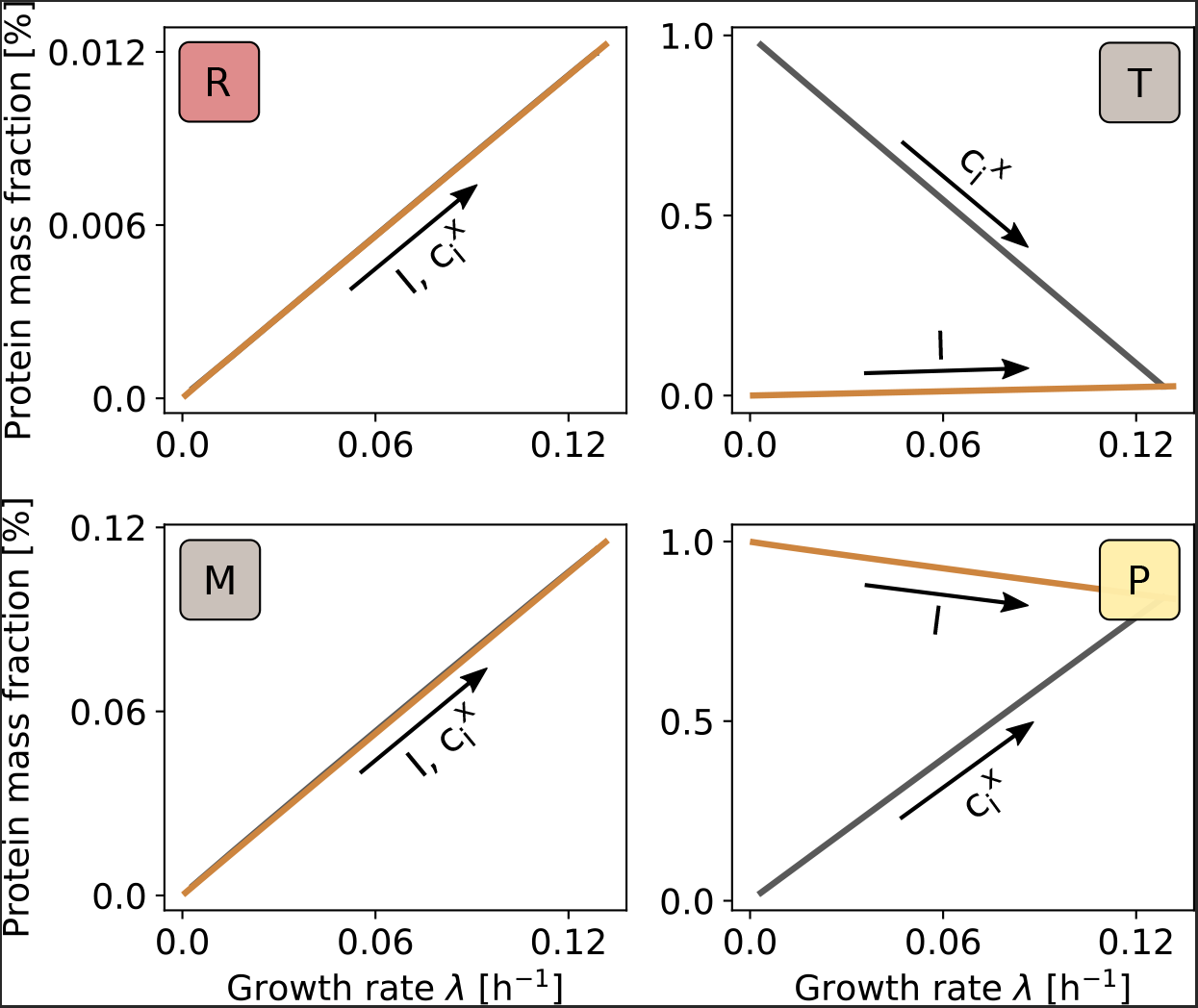
Protein allocation and phototrophic growth laws. Protein mass fractions are linearly dependent on the growth rate. Optimal protein mass fractions are calculated for increasing light intensity *I* and fixed external inorganic carbon 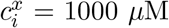, as well as for increasing external inorganic carbon 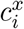 and fixed light intensity *I* = 1000 *μ*E m^-2^ s^-1^. The mass fraction of ribosomal and metabolic proteins (*R, M*) linearly increases with increasing growth rate, irrespective of whether growth is enhanced by increasing 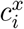 or *I*. The mass fraction of the transporter protein (*T*), however, decreases linearly with increasing carbon availability, whereas it increases for increasing light availability. Vice versa, the mass fraction of the photosynthetic unit (*P*) decreases linearly with increasing light intensity and increases for increasing carbon availability.

All other protein fractions likewise exhibit a linear relationship with respect to the growth rate. The amount of metabolic enzyme *M*, relative to total protein, is linearly increasing with growth rate, irrespective of whether the increase in growth is caused by increasing external inorganic carbon or increasing light intensity. In contrast, the relative mass fraction of the transporter *T* decreases with increasing external inorganic carbon, and increases with increasing light intensity. The former, a decrease of transporter with increasing availability of the carbon source, was also observed in the (heterotrophic) model of Molenaar et al. [10]. The mass fraction of the photosynthetic unit *P* decreases with increasing light intensity and increases with increasing external inorganic carbon. The former agrees with the result of Burnap [4] where the mass fraction of the light-harvesting complex (LHC) decreases with increasing light intensity.

The minimal model shows that bacterial growth laws, as previously determined only for heterotrophic organisms, can be applied to phototrophic growth where the carbon and energy source are separate. In the following, we extend the minimal model to incorporate aspects of photodamage and carbon cycling.

## 4 Beyond the minimal model: photodamage and carbon cycling

### 4.1 High light intensities and photodamage

As yet, the minimal model did not account for two hallmark properties of cyanobacterial phototrophic growth, potential photodamage and carbon cycling. Light absorption damages the photosynthesis machinery proportional to the light intensity [32]. Photodamage has significant impact on the observed growth curve. A number of (mainly phenomenological) minimal models reproduce the decrease in growth rate under high light. In particular, three state models of the photosynthesis-irradiance (PI) curve typically reproduce the inhibitory effect of high light intensities [33]. See Westermark and Steuer [34] for a recent review.

To account for potential photodamage in the minimal model, we follow Han [33] and assume that the active state of the photosynthetic unit *P*\* can be irreversibly damaged by further light absorption. The protein is then degraded into amino acids *aa*, mimicking the repair-cycle of the D1 subunit [32]. The rate of damage (*v_i_*) is assumed to be a linear function of light intensity,

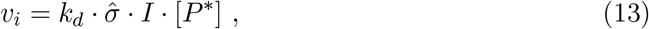

where *k_d_* denotes the first-order rate constant. The inclusion of photodamage modifies the dynamics of the total photosynthetic unit *P*,

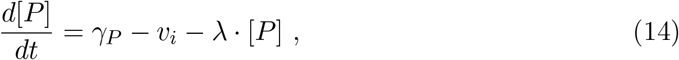

as well as the quasi-steady state expression for the photosynthetic capacity, equation (7),

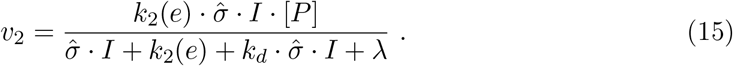

See the supplementary text for the complete set of ODEs. The only additional kinetic parameter is the first-order rate constant *k_d_*. Its value is estimated using experimentally determined growth curves (see section 4.3).

### 4.2 Diffusion and carbon cycling

In addition to potential photodamage, we account for the fact that some forms of inorganic carbon, in particular CO_2_, are permeable. Diffusion of inorganic carbon via the cell membrane results in two effects: at high concentrations of external inorganic carbon, passive diffusion is sufficient to meet the carbon requirements of metabolism, and active transport is not required. Vice versa, for low concentrations of external inorganic carbon and high activity of the transporter reaction, passive diffusion results in leakage of inorganic carbon from the cell. The inclusion of diffusion, together with the active transport mechanism, therefore implements a primitive inorganic carbon concentrating mechanism (CCM): to increase the concentration of intracellular carbon relative to the extracellular concentration, the transport reaction has to operate against a diffusion gradient.

To account for passive diffusion, we assume that the diffusion reaction (*v_d_*) depends on the permeability of the cell membrane *P_m_*, the cell surface *A_cell_* and the gradient between internal and external *c_i_*

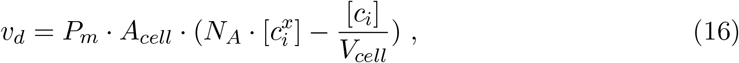

where *V_cell_* and *N_A_* are the cell volume and the Avogadro constant, respectively. For simplicity, *A_cell_* and *V_cell_* are assumed to be constant. The inclusion of the diffusion reaction results in a modified ODE for intracellular inorganic carbon, equation (1),

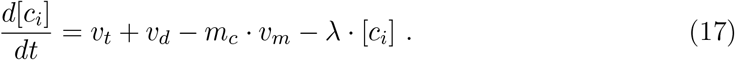

Likewise, the expression for the growth rate, equation (9), is modified (see supplementary text, section 2.4),

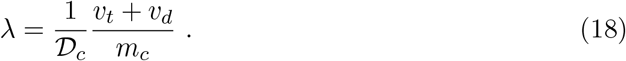

### 4.3 Comparison to experimental growth curves

The inclusion of photodamage and diffusion allows the minimal model to reproduce the characteristic cyanobacterial growth curve. To this end, we measured the growth rate of *Synechocystis* sp. PCC 6803 in a turbidostat culture as a function of light intensity under increased CO_2_, corresponding to carbon-saturated growth. See Materials and Methods for details. The experimental growth curve is shown in figure 4a. The data were used to fit three unknown parameters in the extended model, namely the turnover rate *τ*, the effective absorption cross-section 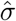, and the rate constant of photodamage *k_d_*. The minimal model and the experimentally derived growth rates are in good agreement (figure 4a).

**Figure 4:**
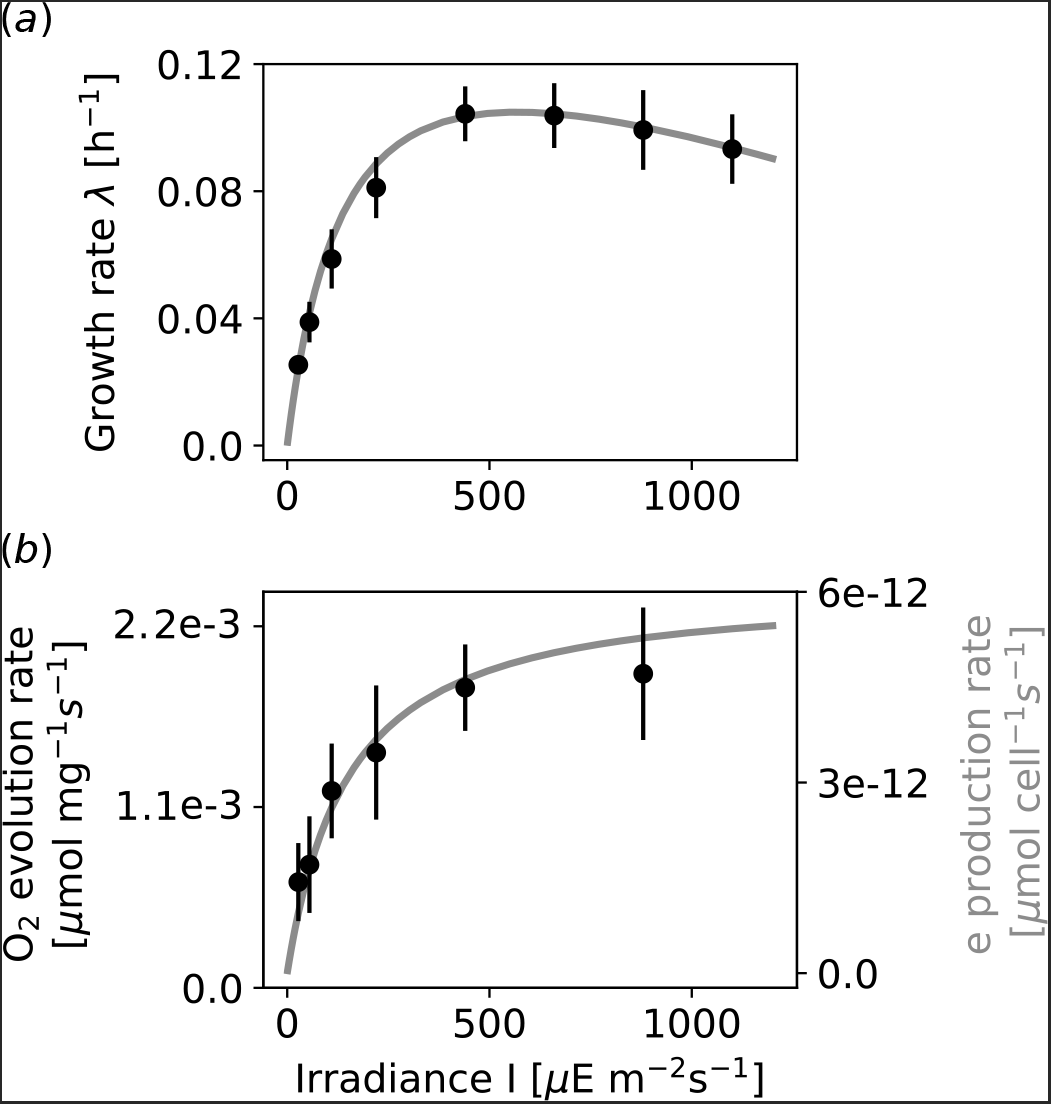
The minimal model reproduces experimental cyanobacterial growth curves. (*a*) The steady-state growth rate of *Synechocystis* sp. PCC 6803 (black circles) was measured experimentally in turbidostat culture for 8 different light intensities with 6 – 11 biological replicates each. Model simulations (solid gray line) were fitted (see section 2.5) using a constant external inorganic carbon value of 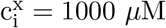 (growth under saturating carbon conditions). (*b*) The model was subsequently used to compare the model-derived photosynthetic turnover rate *v*_2_ (gray line) to the photosynthetic oxygen evolution rate per dry weight. Both curves qualitatively agree (no further parameters were adjusted).

The fitted model can subsequently be used to predict the functional form of the oxygen-evolution rate as a function of light intensity *I*. Since the minimal model does not explicitly account for oxygen evolution, we approximate the oxygen evolution with the turnover of the photosynthetic unit (*v*_2_). No further parameters were adjusted. The functional form of the model-derived turnover rate is in good agreement with the measured oxygen evolution (figure 4b).

### 4.4 Sensitivity analysis and the limits of growth

Using the fitted model, we seek to evaluate the impact of all model parameters, including the availability of external inorganic carbon and light intensity, on the growth rate (sensitivity analysis). To this end, we consider two scenarios: growth under saturating carbon conditions 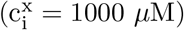, as well as carbon-limited growth 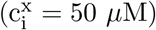. The results are shown in figure 5.

**Figure 5:**
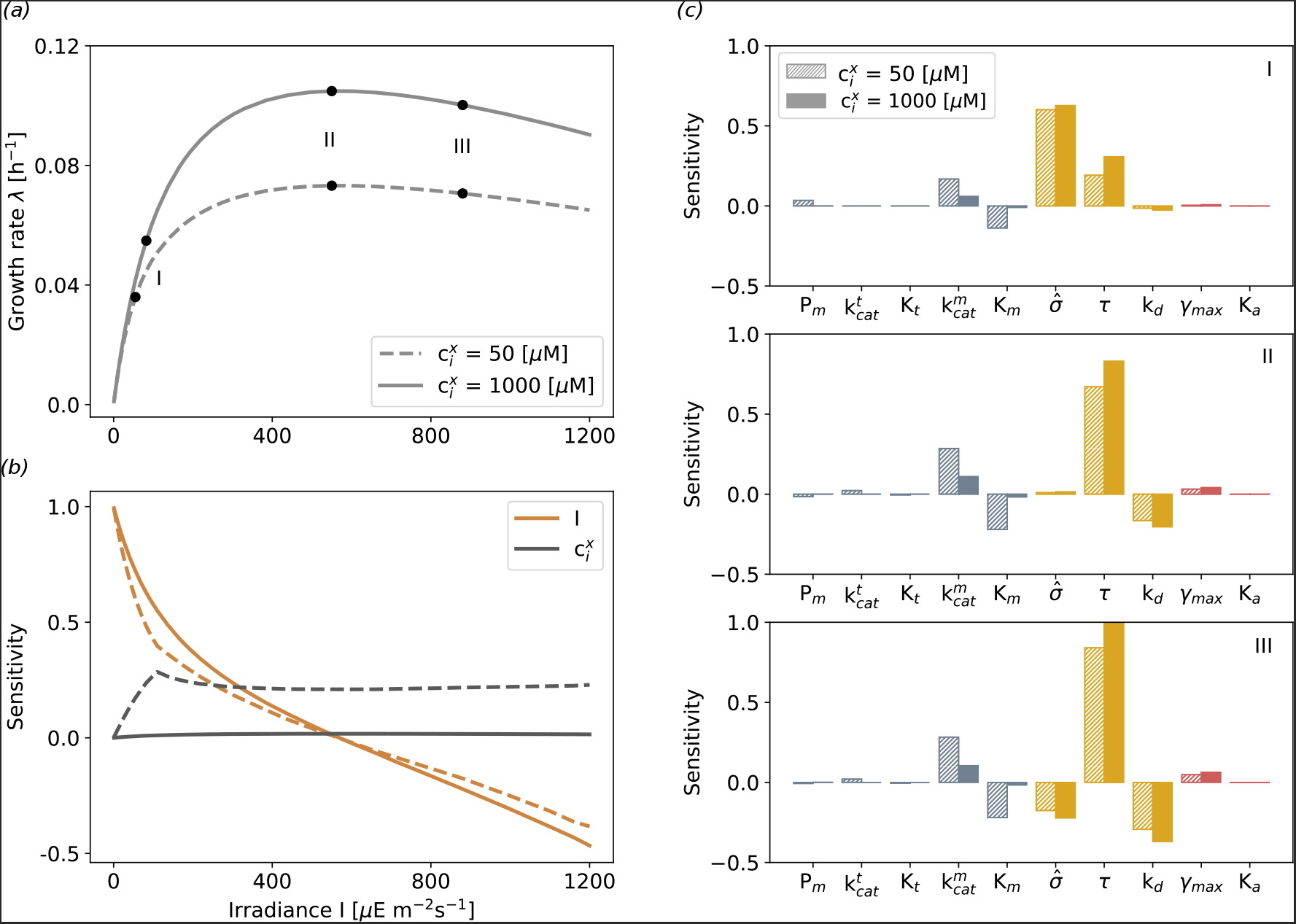
Sensitivity analysis of the extended model with respect to environmental conditions and intracellular parameters. *(a)* Growth curves for two scenarios: carbon-saturating conditions (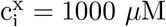, solid line), as well as carbon-limited conditions (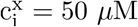, dashed line). *(b)* The (logarithmic) sensitivity with respect to external inorganic carbon 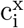 (gray line) and light intensity *I* (orange line). For low light, the sensitivity with respect to light intensity is close to unity, implying an (almost) linear dependence of growth as a function of light intensity The sensitivity with respect to light then decreases until the maximal growth rate is attained. Subsequently, increasing light intensity has a negative impact on the growth rate (photoinhibition). The dependence on extracellular inorganic carbon differs between both growth scenarios. Under carbon-saturating conditions, the sensitivity remains close to zero for the entire growth curve. For low extracellular carbon, the sensitivity increases with increasing light intensity. *(c)* The sensitivity coefficients with respect to intracellular model parameters. The three growth phases correspond to the points marked in panel *(a)*.

The (logarithmic) sensitivity coefficient (see Materials and Methods for a definition) describes the impact of a change in a parameter on the growth rate: a sensitivity coefficient close to unity implies an (approximately) linear dependence. Figure 5b shows the sensitivity coefficient of growth with respect to external parameters *I* and 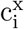. The sensitivity coefficient with respect to intracellular parameters is shown in figure 5c for three different growth phases: (*I*) The first growth phase is characterized by light limitation for both values of external inorganic carbon availability. The model parameter with highest impact on the growth rate is the effective absorption cross-section 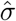 that determines the amount of light absorbed per photosynthetic unit. (II) The second growth phase corresponds to the point where the highest growth rate is attained. In this case, the sensitivity with respect to light intensity is zero. The model parameter with the highest impact on the growth rate is the turnover rate of the photosynthetic unit τ. (III) The third growth phase is characterized by increasing photoinhibition, i.e., a light-induced reduction of growth rate. In this case, the model parameter with the highest impact on the growth rate is the turnover rate of the photosynthetic unit τ, whereas an increasing effective absorption cross-section σ has a negative impact on growth. An increase of the turnover rate τ minimizes the abundance of the photosynthetic unit in its activated state and hence reduces the impact of photoinhibition. Other model parameters, in particular the maximal elongation rate γ_max_, have comparatively low impact on the estimated growth rate.

### 4.5 Effects of photodamage and diffusion on phototrophic growth laws

We are interested in how the incorporation of photodamage and diffusion impacts the phototrophic growth laws shown in figure 3. Figure 6 shows the modified growth laws for the extended model. We consider optimal protein allocation for increasing light intensity for two different amounts of extracellular inorganic carbon (figure 6a), as well as optimal protein allocation for increasing extracellular inorganic carbon for two different light intensities (figure 6b). The corresponding intracellular concentrations of cellular precursors are provided in the supplementary text, figures S6-S7.

**Figure 6:**
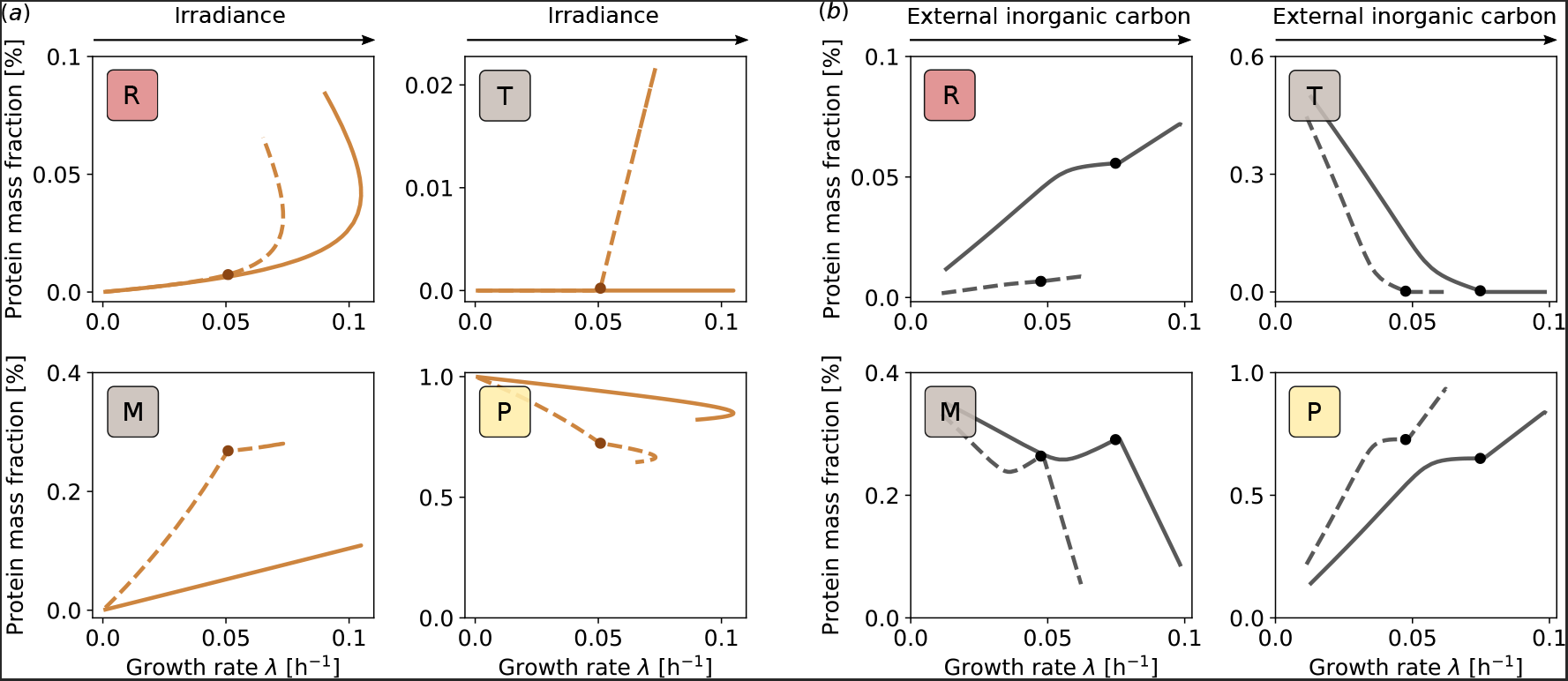
Effects of photodamage and diffusion on phototrophic growth laws. (*a*) Protein mass fractions are displayed for increasing light intensity and fixed external inorganic carbon, 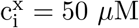 (dashed lines) and 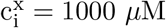 (solid lines). The solid circles indicate the transition point for which the transporter (*T*) mass fraction is more than 0.02%. (*b*) Protein mass fractions for increasing external inorganic carbon and fixed light intensities, I = 100 *μ*E m^-2^ s^-1^ (dashed lines) and I = 1000 *μ*E m^-2^ s^-1^ (solid lines). The solid circles indicate the transition point for which the transporter mass fraction is less than 0.3%.

For increasing light intensity *I* (and fixed 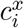) the ribosomal mass fraction first increases proportional to the growth rate. For high light intensities and fast growth, however, the ribosomal mass continues to increase even as the growth rate again decreases due to photoinhibition. A similar phenomenon is observed for *E. coli* when nutrient amount is fixed and translational inhibitors are added to the medium [17]: the cell compensates the inhibition of ribosomes by increasing the ribosomal mass fraction. In our case, the light-induced repair-cycle of the photosynthetic unit acts analogously to a translational inhibitor and requires to increase the ribosomal mass fraction. We note that, as a function of light intensity, the ribosomal mass fraction retains its linear dependence (see supplementary text, figure S7b).

For increasing light intensity *I* (and fixed 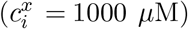) the amount of transporter protein *T* exhibits a switch-like behavior. For high availability of external inorganic carbon 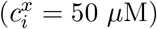, passive diffusion is sufficient to meet the carbon requirements of the cell and the transporter protein is not expressed. For low external inorganic carbon 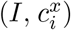, passive diffusion is only sufficient up to a critical threshold in the growth rate after which the transporter protein *T* is expressed (see supplementary text, figure S5). Conditional expression of bicarbonate transporter has been reported in the literature [35, 36].

The remaining growth laws are qualitatively similar to the results shown in figure 3 - albeit typically with modifications to the simple linear relationships. In particular, the switch between passive diffusion and active transport, indicated with a dot in figure 6, induces shifts in the optimal protein allocation. Another noteworthy difference between the initial model and the extended model is the strong increase in the mass fraction of the metabolic protein *M*. Due to the inclusion of passive diffusion, the intracellular abundance of inorganic carbon is significantly lower in the extended model (up to two orders of magnitude). Due to the low affinity of the metabolic protein, the protein is not fully saturated for low external inorganic carbon - a fact that must be compensated by an increased mass fraction of the metabolic protein.

Overall, the results shown in figure 6 indicate that the growth laws for phototrophic growth are qualitatively retained when photodamage and diffusion are considered within the model - albeit with several important modifications. We conjecture that the actual protein mass fractions of phototrophic growth may be more complex than the simple linear dependencies hitherto described for heterotrophic growth.

## 5 Discussion and Conclusions

Cyanobacteria are promising host organisms for green biotechnology. However, our understanding of the molecular limits of cyanobacterial growth and the associated phototrophic growth laws is insufficient. Here, we developed a minimal kinetic model of cyanobacterial phototrophic growth to describe optimal protein allocation under different environmental conditions. The model incorporates key processes related to phototrophic growth, i.e., carbon uptake, metabolism, photosynthesis and protein translation. Different from corresponding models of heterotrophic growth, we distinguish between two potentially limiting nutrients, external inorganic carbon and light.

The model is able to reproduce experimentally obtained phototrophic growth curves, including the inhibition of growth at high light intensities. The minimal model also reproduces typical growth laws known from heterotrophic bacteria, in particular the fact that ribosomal mass fraction increases linearly with increasing growth rate (irrespective of whether the increase in growth rate is brought about by an increase in light intensity or an increase in availability of inorganic carbon). The growth laws obtained here are in good agreement with the effects of light intensity on proteome allocation modeled by Burnap [4]: the mass fraction of the photosynthetic unit (and hence light-harvesting complexes) decreases with increasing light intensity. Vice versa, the mass fraction of metabolic enzymes increases. The inclusion of potential photodamage and diffusion results in more complex phototrophic growth laws, while general properties and trends are preserved. In particular photodamage, mimicking the photoinhibition repair-cycle of enzymatic degradation and synthesis of the D1 protein, places an additional burden on ribosomal capacity - the resulting curves are similar to growth laws obtained for heterotrophic bacteria that are subject to a translational inhibitor. We conjecture, however, that the minimal model, where photodamage primarily increases the requirement of translation, overestimates the effects of photodamage on the ribosomal mass fraction.

Compared to heterotrophic organisms, quantitative data on cyanobacterial growth laws under different growth conditions are scarce. A few experimental studies have considered carbon and phosphorus allocation [37, 38], as well as rRNA content [39, 40]. The latter point to a linear increase for intermediate growth rates, in good agreement with modeling results. The amount of rRNA per cell, however, plateaus or decreases at highest growth rates (explained at least in part by a decrease in cell size). Clarification would require direct measurements of cell size and protein content [40]. Likewise, rRNA content is constant at lower growth rates [40]. A similar deviation from model-derived optimality has already been observed for *E. coli* [14] - here, the higher relative amount of the ribosomal mass fraction at low growth rates is likely the result of an anticipatory adaptation to potential higher growth rates when nutrient availability increases.

The data that allow the most direct comparison with experimental protein allocation is the recent study of Bernstein et al. [6] - measuring transcriptional responses to changing irradiance, and hence changing growth rate, in a turbidostat. Among the genes whose relative abundance of transcripts increases in direct proportion with growth rate were genes involved in translation (ribosomal proteins), amino acid biosynthesis and other genes involved in central carbon metabolism [6]. Among the genes whose relative abundance of transcripts inversely correlates with growth rate were genes encoding photosystems I and II antenna proteins, as well as transcripts related to niche-adaptive protein functions [6]. These results indicate similar trends as predicted here. For a detailed understanding of protein optimal allocation, however, further quantitative studies are needed. Computational models, such as the one presented here, will undoubtedly play an important role in interpreting such quantitative studies with respect to optimal adaptation of cyanobacteria to different growth conditions.

While the current minimal model captures key properties of phototrophic growth, the model may be extended to address specific cellular functions in more details. For example, currently, we only consider a minimal CCM and neglect photorespiration, as well as other photoprotective mechanisms. Future work may include a more detailed representation of the electron transport chain, differentiating between the provision of energy and reductants, as well as between linear and cyclic electron transport. Growth conditions may include fluctuating light to study energy dissipation and adaptation to varying light intensities. An ultimate goal is to improve our understanding of day/night regulation in diurnal environments, necessitating the inclusion of storage metabolism. Furthermore, we also envision applications of this and similar minimal growth models to bridge the gap between molecular physiology and ecology. Indeed, it has recently been argued that current plankton models should place more emphasis on the underlying mechanisms of cell physiology, rather than on empirical parameter fitting, to allow trade-offs between resource allocation and ecological dynamics as emergent properties [41]. The model proposed here recapitulates known growth laws but connects them to underlying molecular mechanisms - and therefore may serve as a starting point to integrate ecophysiology with systems biology.

## 6 Materials and Methods

### 6.1 Experimental procedures and growth data

As a reference organism, we used *Synechocystis* sp. PCC 6803, substrain GT-L (*Synechocystis* hereafter). *Synechocystis* was cultivated in a flat panel photobioreactor [42] under 27.5−1100 *μ*mol photons m^-2^ s^-1^ of red light (λ_max_ ≈ 633 nm, λ 1/2 ≈ 20 nm, Luxeon LXHLPD09, Future Lighting Solutions, Montreal, QC, Canada) supplemented with low portion of blue light (25 *μ*mol photons m^-2^ s^-1^, λ_max_ ≈ 445 nm, λ1/2 ≈ 20 nm, Luxeon LXHL-PR09; Future Lighting Solutions). The cultures were cultivated at 32°C and the culture suspensions were bubbled by air supplemented with 0.5% CO_2_ (v/v). The cultures were cultivated in a quasi-continuous regime operated as turbidostat according to [43]. Briefly, the exponentially growing cultures were periodically diluted with fresh culture medium. The dilution was based on automatic measurement of culture optical density at 680 nm (OD_680_); the OD_680_ range was set to 0.52 - 0.58 (approximately 10^7^ cells ml^-1^). The cultures were cultivated under each red light intensity for at least 24 hours. This period was long enough to reach growth stability, i.e. to adapt to particular light conditions. After the cultures reached growth stability, samples were taken for measurement of cell count, cellular dry weight and concentration on chlorophyll a in the *Synechocystis* cells. Specific growth rates were calculated from changes in OD_680_ by applying the exponential regression model.

Dissolved oxygen was monitored by InPro6800 electrodes (Mettler-Toledo, Inc., Columbus, OH, USA). The oxygen evolution/respiration measurements were performed in the photobioreactor cuvette by turning off the cultures aeration for 10 min, through a 5 min light period and a 5 min dark period. Oxygen evolution/respiration rates were normalized per chlorophyll *a* content in *Synechocystis* cells, determined according to [44]. The cell count was measured with a Cellometer Auto M10 (Nexcelom Bioscience, Lawrence, MA, USA). The dry weight was measured with analytical balances XA105DR (Mettler-Toledo, Greifensee, CH).

### 6.2 Model implementation

The complete set of ordinary differential equations (ODEs) and their parametrization is given in the supplementary text. The ODE system is implemented in Python 2.7 and available under the name *minimal_model.py*. To simulate the model results, we used the method odeint from the python package *scipy.integrate* for solving the ODEs and subsequently the method *minimize* from *scipy.optimize* for solving the optimization problem. The script to execute both steps is given in *simulate.py*.

### 6.3 Sensitivity analysis

The sensitivity is estimated for all model parameters and external conditions 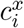, except for the ribosome fractions (*β*). The (logarithmic) sensitivity s_i_ of the growth rate λ for a given parameter *p_i_* is by the derivative

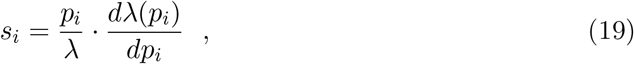

and approximated by finite-size difference (0.1%) of a parameter in steady state at the optimized growth rate. The value of the (logarithmic) sensitivity *s_i_* corresponds to the kinetic order, i.e., a value close to one indicates a linear dependence of the growth rate on the parameter *p_i_*.

## Data accessibility

Python code of the model and for simulating the optimization results, as well as the experimental data, are given in the electronic supplementary material.

## Author’s contributions

R.S. conceived of the study and designed the study; M.F. carried out all computational simulations and drafted the manuscript; T.Z., C.L., and J.C. carried out the growth experiments and carried out data analyses; M.F. and R.S. wrote the manuscript. All authors gave final approval for publication.

## References

[1] Daniel C Ducat, Jeffrey C Way, and Pamela A Silver. Engineering cyanobacteria to generate high-value products. Trends Biotechnol., 29(2):95–103, 2011. doi: 10.1016/j.tibtech.2010.12.003.

[2] Tomáš Zavřel, Jan Červený, Henning Knoop, and Ralf Steuer. Optimizing cyanobacterial product synthesis: Meeting the challenges. Bioengineered, 7:490–496, 2016. ISSN 2165-5987. doi: 10.1080/21655979.2016.1207017.

[3] Aharon Oren. Cyanobacteria An Economic Perspective - Cyanobacteria: biology, ecology and evolution, chapter 1, pages 12–13. John Wiley & Sons, 2014.

[4] Robert L Burnap. Systems and photosystems: cellular limits of autotrophic productivity in cyanobacteria. Front. Bioeng. Biotechnol., 3:1, 2015. doi: 10.3389/fbioe.2015.00001.

[5] Jingjie Yu, Michelle Liberton, Paul F Cliften, Richard D Head, Jon M Jacobs, Richard D Smith, David W Koppenaal, Jerry J Brand, and Himadri B Pakrasi. Synechococcus elongatus UTEX 2973: a fast growing cyanobacterial chassis for biosynthesis using light and CO2. Sci. Rep., 5:8132, 2015. ISSN 2045-2322. doi: 10.1038/srep08132.

[6] Hans C Bernstein, Ryan S McClure, Eric A Hill, Lye Meng Markillie, William B Chrisler, Margie F Romine, Jason E McDermott, Matthew C Posewitz, Donald A Bryant, Allan E Konopka, James K Fredrickson, and Alexander S Beliaev. Unlocking the constraints of cyanobacterial productivity: Acclimations enabling ultrafast growth. mBio, 7, 2016. ISSN 2150-7511. doi: 10.1128/mBio.00949-16.

[7] Marco Rügen, Alexander Bockmayr, and Ralf Steuer. Elucidating temporal resource allocation and diurnal dynamics in phototrophic metabolism using conditional FBA. Sci. Rep., 5:15247, 2015. doi: 10.1038/srep15247.

[8] Thomas J Mueller, Justin L Ungerer, Himadri B Pakrasi, and Costas D Maranas. Identifying the metabolic differences of a fast-growth phenotype in Synechococcus UTEX 2973. Sci. Rep., 7:41569, 2017. ISSN 2045-2322. doi: 10.1038/srep41569.

[9] Alexandra-M. Reimers, Henning Knoop, Alexander Bockmayr, and Ralf Steuer. Cellular trade-offs and optimal resource allocation during cyanobacterial diurnal growth. Proc. Natl. Acad. Sci. U.S.A., 114(31):E6457–E6465, 2017. doi: 10.1073/pnas.1617508114.

[10] Douwe Molenaar, Rogier van Berlo, Dick de Ridder, and Bas Teusink. Shifts in growth strategies reflect tradeoffs in cellular economics. Mol. Syst. Biol., 5:323, 2009. doi: 10.1038/msb.2009.82.

[11] Jacques Monod. The growth of bacterial cultures. Annu. Rev. Microbiol., 3:371–394, 1949. doi: 10.1016/b978-0-12-460482-7.50020-8.

[12] Moselio Schaechter. A brief history of bacterial growth physiology. Front. Microbiol., 6:289, 2015. ISSN 1664-302X. doi: 10.3389/fmicb.2015.00289.

[13] F C Neidhardt. Bacterial growth: constant obsession with dn/dt. J. Bacteriol., 181: 7405–7408, 1999. ISSN 0021-9193.

[14] Allen G Marr. Growth rate of Escherichia coli. Microbiol. Rev., 55(2):316–333, 1991.

[15] R E Ecker and M Schaechter. Ribosome content and the rate of growth of Salmonella typhimurium. Biochim. Biophys. Acta, 76:275–279, 1963. ISSN 0006-3002. doi: 10.1016/0926-6550(63)90040-9.

[16] Stefan Klumpp, Zhongge Zhang, and Terence Hwa. Growth rate-dependent global effects on gene expression in bacteria. Cell, 139:1366–1375, 2009. doi: 10.1016/j.cell.2009.12.001.

[17] Matthew Scott, Carl W. Gunderson, Eduard M. Mateescu, Zhongge Zhang, and Terence Hwa. Interdependence of cell growth and gene expression: Origins and consequences. Science, 330:1099–1102, 2010. doi: 10.1126/science.1192588.

[18] Evert Bosdriesz, Douwe Molenaar, Bas Teusink, and Frank J Bruggeman. How fast-growing bacteria robustly tune their ribosome concentration to approximate growth-rate maximization. FEBS J., 282:2029–2044, 2015. doi: 10.1111/febs.13258.

[19] Arijit Maitra and Ken A Dill. Bacterial growth laws reflect the evolutionary importance of energy efficiency. Proc. Natl. Acad. Sci. U.S.A., 112(2):406–411, 2015. doi: 10.1073/pnas.1421138111.

[20] Andrea Y. Weiβe, Diego A. Oyarzún, Vincent Danos, and Peter S. Swain. Mechanistic links between cellular trade-offs, gene expression, and growth. Proc. Natl. Acad. Sci. U.S.A., 112(9):E1038–47, 2015. doi: 10.1101/014787.

[21] R O Megard D W Tonkyn, and W H Senft. Kinetics of oxygenic photosynthesis in planktonic algae. J. Plankton Res., 6(2):325–337, 1984. doi: 10.1093/plankt/6.2.325.

[22] P H C Eilers and J C H Peeter. A model for the relationship between light intensity and the rate of photosynthesis in phytoplankton. Ecol. Modell., 42(3-4):199–215, 1988. doi: 10.1016/0304-3800(88)90057-9.

[23] C Zonneveld. Modeling effects of photoadaption on the photosynthesis-irradiance curve. J. Theor. Biol., 186(3):381–388, 1997. doi: 10.1006/jtbi.1997.0400.

[24] Bo-Ping Han. Photosynthesis-irradiance response at physiological level: a mechanistic model. J. Theor. Biol., 213(2):121–127, 2001. doi: 10.1006/jtbi.2001.2413.

[25] Hans Bremer and Patrick P Dennis. Modulation of chemical composition and other parameters of the cell at different exponential growth rates. EcoSal Plus, 3(1), 2008. doi: 10.1128/ecosal.5.2.3.

[26] Henning Knoop, Marianne Gründel, Yvonne Zilliges, Robert Lehmann, Sabrina Hoffmann, Wolfgang Lockau, and Ralf Steuer. Flux balance analysis of cyanobacterial metabolism: The metabolic network of Synechocystis sp. PCC 6803. PLoS Comput. Biol., 9(6):e1003081, 2013. doi: 10.1371/journal.pcbi.1003081.

[27] Yehouda Marcus, Hagit Altman-Gueta, Aliza Finkler, and Michael Gurevitz. Mutagenesis at two distinct phosphate-binding sites unravels their differential roles in regulation of Rubisco activation and catalysis. J. Bacteriol., 187(12):222–4228, 2005. doi: 10.1128/jb.187.12.4222-4228.2005.

[28] Niall M Mangan and Michael P Brenner. Systems analysis of the CO2 concentrating mechanism in cyanobacteria. eLife, 3:e02043, 2014. doi: 10.7554/eLife.02043.

[29] Klaus Dornmair, Peter Overath, and Fritz Jähnig. Fast measurement of galactoside transport by lactose permease. J. Biol. Chem., 26(1):342–346, 1989.

[30] Tatsuo Omata, Yukari Takahashi, Osamu Yamaguchi, and Takashi Nishimura. Structure, function and regulation of the cyanobacterial high-affinity bicarbonate transporter, BCT1. Funct. Plant Biol., 29(3):151–159, 2002.

[31] Mak A Saito, Tyler J Goepfert, and Jason T Ritt. Some thoughts on the concept of colimitation: Three definitions and the importance of bioavailability. Limnol. Oceanogr., 53(1):276–290, 2008. doi: 10.4319/lo.2008.53.1.0276.

[32] Esa Tyystjärvi. Photoinhibition of Photosystem ii and photodamage of the oxygen evolving manganese cluster. Coord. Chem. Rev., 252(3-4):361–376, 2008. doi: 10.1016/j.ccr.2007.08.021.

[33] Bo-Ping Han. A mechanistic model of algal photoinhibition induced by photodamage to photosystem-ii. J. Theor. Biol., 214(4):519–527, 2002. doi: 10.1006/jtbi.2001.2468.

[34] Stefanie Westermark and Ralf Steuer. Toward multiscale models of cyanobacterial growth: A modular approach. Front. Bioeng. Biotechnol., 4:95, 2016. doi: 10.3389/fbioe.2016.00095.

[35] P J McGinn G D Price, and M R Badger. High light enhances the expression of low-CO2-inducible transcripts involved in the CO2-concentrating mechanism in Synechocystis sp. PCC6803. Plant Cell Environ., 27(5):615–626, 2004. doi: 10.1111/j.1365-3040.2004.01175.x.

[36] Robert L Burnap, Martin Hagemann, and Aaron Kaplan. Regulation of CO2 concentrating mechanism in cyanobacteria. Life (Basel), 5:348–371, 2015. ISSN 2075-1729. doi: 10.3390/life5010348.

[37] Tyler D B MacKenzie, Jeanette M Johnson, Amanda M Cockshutt, Robert A Burns, and Douglas A Campbell. Large reallocations of carbon, nitrogen, and photosynthetic reductant among phycobilisomes, photosystems, and Rubisco during light acclimation in Synechococcus elongatus strain PCC7942 are constrained in cells under low environmental inorganic carbon. Arch. Microbiol., 183:190–202, 2005. ISSN 0302-8933. doi: 10.1007/s00203-005-0761-1.

[38] John A Raven. RNA function and phosphorus use by photosynthetic organisms. Front. Plant Sci., 4:536, December 2013. ISSN 1664-462X. doi: 10.3389/fpls.2013.00536.

[39] Brian J Binder and Ying Chun Liu. Growth rate regulation of rRNA content of a marine Synechococcus (cyanobacterium) strain. Appl. Environ. Microbiol., 64(9): 3346–3351, 1998.

[40] Alexandra Z Worden and Brian J Binder. Growth regulation of rRNA content in Prochlorococcus and Synechococcus (marine cyanobacteria) measured by whole-cell hybridization of rRNA-targeted peptide nucleic acids. J. Phycol., 39(3):527–534, 2003. doi: 10.1046/j.1529-8817.2003.01248.x.

[41] J I Allen and L Polimene. Linking physiology to ecology: towards a new generation of plankton models. J. Plankton Res., 33(7):989–997, 2011. doi: 10.1093/plankt/fbr032.

[42] Ladislav Nedbal, Martin Trtílek, Jan Cervený, Ondrej Komárek, and Himadri B Pakrasi. A photobioreactor system for precision cultivation of photoautotrophic microorganisms and for high-content analysis of suspension dynamics. Biotechnol. Bioeng., 100:902–910, 2008. ISSN 1097-0290. doi: 10.1002/bit.21833.

[43] Tomáš Zavřel, Maria A Sinetova, Diana Búzová, Petra Literáková, and Jan Červeny. Characterization of a model cyanobacterium Synechocystis sp. PCC 6803 autotrophic growth in a flat-panel photobioreactor. Eng. Life Sci., 15(1):122–132, 2015. doi: doi:10.1002/elsc.201300165.

[44] T Zavřel, MA Sinetova, and J Červený. Measurement of chlorophyll a and carotenoids concentration in cyanobacteria. Bio Protoc., 5:1–5, 2015. URLhttp://www.bio-protocol.org/e1467.

